# Interrupted reprogramming into induced pluripotent stem cells does not rejuvenate human mesenchymal stromal cells

**DOI:** 10.1101/344903

**Authors:** Carolin Göbel, Roman Goetzke, Thomas Eggermann, Wolfgang Wagner

## Abstract

Replicative senescence hampers application of mesenchymal stromal cells (MSCs) because it limits culture expansion, impairs differentiation potential, and hinders reliable standardization of cell products. MSCs can be rejuvenated by reprogramming into induced pluripotent stem cells (iPSCs), which is associated with complete erasure of age- and senescence-associated DNA methylation (DNAm) patterns. However, this process is also associated with erasure of cell-type and tissue-specific epigenetic characteristics that are not recapitulated upon re-differentiation towards MSCs. In this study, we therefore followed the hypothesis that overexpression of pluripotency factors under culture conditions that do not allow full reprogramming might reset senescence-associated changes without entering a pluripotent state. MSCs were transfected with episomal plasmids and either successfully reprogrammed into iPSCs or cultured in different media with continuous passaging every week. Overexpression of pluripotency factors without reprogramming did neither prolong culture expansion nor ameliorate molecular and epigenetic hallmarks of senescence. Notably, transfection resulted in immortalization of one cell preparation with gain of large parts of the long arm of chromosome 1. Taken together, premature termination of reprogramming does not result in rejuvenation of MSCs and harbours the risk of transformation. This approach is therefore not suitable to rejuvenate cells for cellular therapy.

## Introduction

Mesenchymal stromal cells (MSCs) raise high expectations for cellular therapy and tissue engineering, particularly due to ease of their isolation^1^. However, application of MSCs is hampered by functional changes caused by replicative senescence during culture expansion^2^. The derivation of MSCs from induced pluripotent stem cells (iPSCs) may help to overcome at least some of these limitations^3,4^. iPSCs can be expanded infinitively without any signs of replicative senescence. Subsequently, iPSC-derived MSCs (iMSCs) can be generated under standardized conditions to provide an unlimited source of younger and more homogeneous cell preparations. In fact, iMSCs reveal similar morphology, surface markers, gene expression profiles, and *in vitro* differentiation potential as primary MSCs^3^. Despite these similarities, iMSCs remain molecularly distinct from primary MSCs, which might be attributed to erasure of epigenetic characteristics of cell type and tissue by conversion into iPSCs^3^. Furthermore, their state of cellular aging, such as senescence-associated epigenetic modifications, seems to be reset in iPSCs and gradually reacquired while differentiating towards MSCs^3^^-^^5^.

Reprogramming of cells into iPSCs is usually achieved by overexpression of pluripotency factors, resulting in a ground state similar to embryonic stem cells (ESCs)^6^. This process seems to be directly associated with rejuvenation with regard to various molecular markers: Expression of senescence-associated genes^7^, telomere lengths^8^, age-associated DNA methylation^3^, and mitochondrial activity^7^ are reset upon reprogramming. However, full cellular reprogramming is also accompanied by complete dedifferentiation and by a risk of teratoma formation *in vivo*^9^. Therefore, it has been suggested that somatic cells can be rejuvenated by partial reprogramming. Ocampo et al. recently demonstrated that hallmarks of aging can be ameliorated through partial reprogramming by cyclic induction of OSKM factors (OCT4, SOX2, KLF4, C-MYC) in prematurely aged mice^10^. Furthermore, Guo et al. used a similar approach for doxycycline-inducible short-term induction of OSKM factors in murine epithelial cells and generated large numbers of progenitor-like cells *in vitro*^11^. In both cases, OSKM induction was interrupted before the cells entered pluripotency.

In this study, we therefore followed the hypothesis that overexpression of reprogramming factors in human MSCs, without full conversion into a pluripotent state, might reverse cellular aging without resetting their functional properties.

## Results

### Overexpression of reprogramming factors without full reprogramming did not support culture expansion of MSCs

Mesenchymal stromal cells of three different donors were transfected at passage six with episomal plasmids coding for OSKM factors, LIN28, and siRNA for P53, which provides an efficient method for integration-free reprogramming into iPSCs^12^. In addition, transfected cells could be detected by expression of the enhanced green fluorescent protein (EGFP) (Fig. 1a,b). These transfected MSCs were successfully reprogrammed into iPSCs when seeded onto mouse embryonic feeder cells (MEFs) with standard iPSC culture conditions (Suppl. Fig. S1a). The pluripotent state of these iPSCs was further confirmed by Epi-Pluri-Score analysis^13^ (Suppl. Fig. S1b). In contrast, transfected MSCs that were maintained in MSC-culture medium comprising human platelet lysate (hPL) did not acquire iPSC morphology and entered replicative senescence after 42 days as also observed for the non-transfected controls (Fig. 1c).

**Figure 1:**
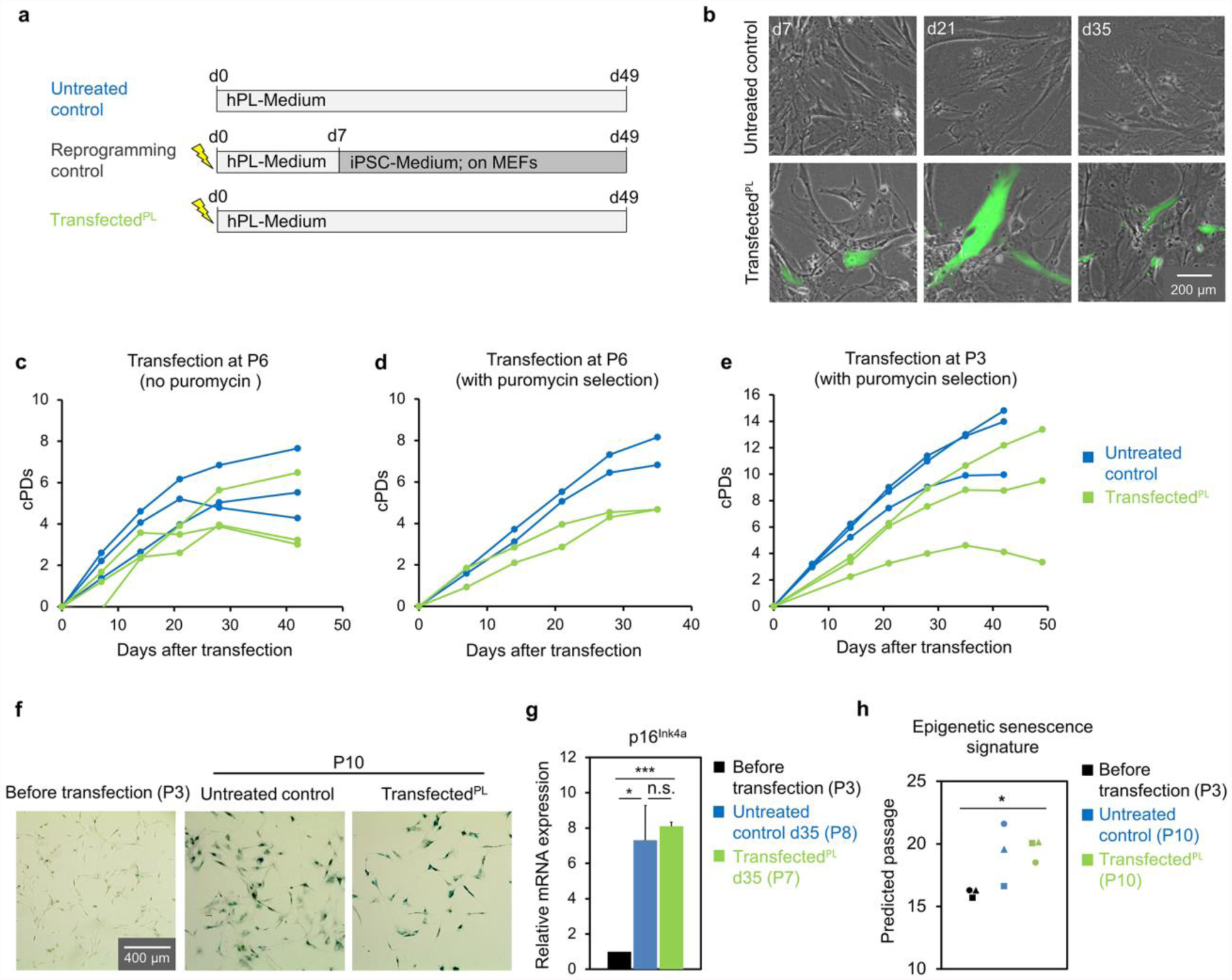
Overexpression of pluripotency factors did not counteract replicative senescence. (**a**) Schematic presentation of culture conditions (hPL = human platelet lysate; the lightning symbol exemplifies electroporation with reprogramming factors; MEFs = mouse embryonic fibroblast feeder layer). (**b**) Fluorescence microscopic analysis demonstrates presence of some EGFP-positive transfected cells throughout culture expansion. (**c-e**) Cumulative population doublings (cPDs) after transfection of MSCs: (**c**) at passage 6 without puromycin selection; (**d**) at passage 6 with puromycin selection; and (**e**) at passage 3 with puromycin selection. (**f**) Exemplary bright-field images of stainings of senescence-associated beta-galactosidase. (**g**) Relative mRNA expression of p16^Ink4a^ as evaluated by qRT-PCR (n = 3, mean ± SD, two-tailed paired Student’s t-test, *p ≤ 0.05, ***p ≤ 0.001). (**h**) Comparison of real and predicted passage numbers for transfected and untransfected MSCs with the Epigenetic-Senescence-Signature^18^ (n = 3, *p ≤ 0.05).

We reasoned that the rejuvenating effect might be concealed by the majority of non-transfected cells. Therefore, we repeated the experiments with addition of a puromycin resistance gene to the transfection and subsequent puromycin selection (1 µg/mL for two days). The results were very similar without evidence that transfected cells with hPL-medium reached higher cumulative population doublings (cPDs) as compared to non-transfected controls (without selection; Fig. 1d). Replicative senescence was shown to be a barrier for reprogramming into iPSCs^7^, and thus we transfected MSC preparations at an earlier passage (P3) in the next experiment. In fact, the initial transfection efficiency was higher at passage three as compared to passage six (Suppl. Fig. S1c). However, there was no beneficial effect on long-term growth curves either (Fig. 1e).

### Molecular markers of cellular senescence were not affected by overexpression of pluripotency factors

Although overexpression of pluripotency factors in cells that were maintained in hPL-medium did not impact on culture expansion, we aimed for a better characterization. The cells revealed typical MSC-like immunophenotype and osteogenic, adipogenic, and chondrogenic differentiation potential (Suppl. Fig. S2). Staining for senescence-associated beta-galactosidase (SA-ß-gal) did not show clear differences between transfected cells and untreated controls (Fig. 1f). Furthermore, expression of p16^Ink4a^, which is indicative for cellular senescence^14,15^, was significantly upregulated in MSCs of later passage as compared to early passage (untreated control: p = 0.03, transfected^PL^: p = 0.0004), whereas there was no difference between transfected cells and untreated controls (Fig. 1g). We have previously demonstrated that DNA methylation levels at specific CG dinucleotides (CpG sites) reveal highly reproducible and continuous changes during culture expansion^16^^-^^18^. Hence, DNA methylation analysis at only six CpG sites provides an Epigenetic-Senescence-Signature that can be used as a surrogate marker to estimate the number of passages^17,18^. Although our Epigenetic-Senescence-Signature on untreated and transfected MSCs generally overestimated passage numbers, passage predictions clearly increased between early and late passage (transfected^PL^: p = 0.03) while there was no effect resulting from the overexpression of pluripotency factors (Fig. 1h).

### Alternative culture conditions for rejuvenation of MSCs with intermittent use of iPSC-culture medium

Since the hPL-culture medium might already interfere with initial reprogramming steps, which may be relevant for rejuvenation, we alternatively tested various culture conditions in parallel (Fig. 2a): i) Untransfected cells were cultured in standard conditions on plastic with hPL-medium (untreated control). Transfected cells were transferred on day 14 either ii) on MEFs in iPSC-medium to achieve successful reprogramming (reprogramming control); iii) in hPL-medium (PL) without MEFs, as described before; iv) in iPSC-medium with regular passaging on tissue culture plastic (PL-i); v) in iPSC-medium for 14 days before changing back to hPL-medium (PL-i-PL), or vi) on MEFs with iPSC-medium for 21 days with passaging every week before changing back to hPL-medium (PL-i(MEF)-PL). All experiments were performed with MSC preparations of three different donors at P3 and puromycin selection (1.5 µg/mL for three days). None of the culture conditions, apart from the reprogramming controls, consistently resulted in extended culture expansion or a higher number of cPDs (Fig. 2b). In fact, culture expansion was significantly impaired by the transfection and selection procedures. Staining for SA-ß-gal at day 42 appeared to be lower in all transfected conditions as compared to untreated controls, which might be attributed to the reduced number of cPDs (Fig. 2c). The Epigenetic-Senescence-Signature did not provide evidence of rejuvenation for any transfected conditions, apart from reprogramming controls that resulted in iPSCs that were estimated close to zero passages (Suppl. Fig. S3). Taken together, none of the culture conditions tested resulted in consistent rejuvenation of MSCs. However, MSCs of donor 2 seemed to be immortalized in culture conditions PL, PL-I, and PL-i-PL – and these cells were then further characterized.

**Figure 2:**
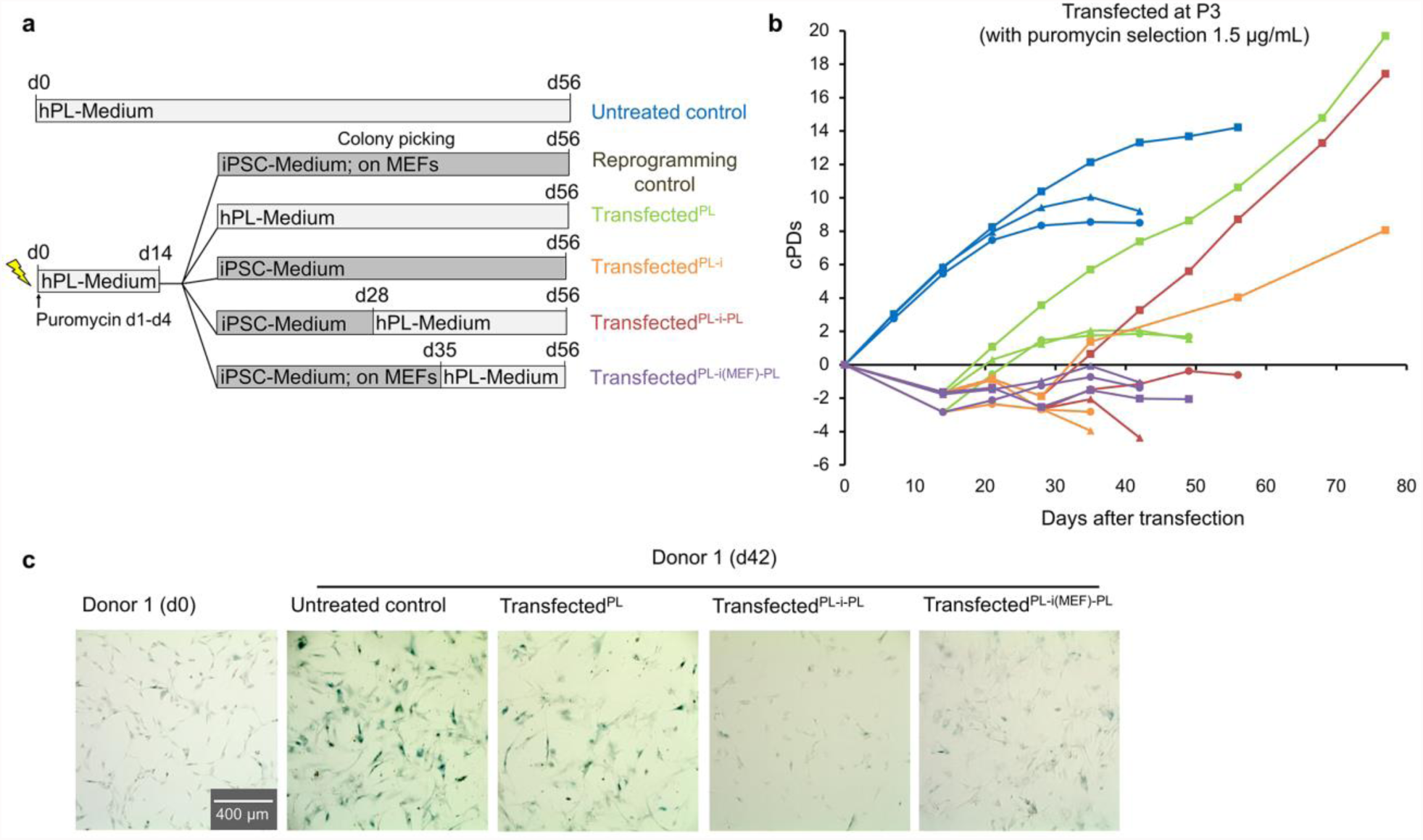
Interrupted reprogramming of MSCs with various culture regimen. (**a**) Schematic presentation of various culture conditions that were compared in parallel. (**b**) Cumulative population doublings (cPDs) after transfection. (**c**) Exemplary images of senescence-associated beta-galactosidase stainings after transfection (d0) and culture expansion (day 42).

### Transfection resulted in a subclone with genetic abnormalities

The immortalized cell preparations of donor 2 revealed typical MSC morphology until day 49 and then gradually changed into smaller cells with epithelial-like morphologies (Fig. 3a). This was accompanied by a loss of surface marker expression for CD73 and CD105 (Fig. 3b). Moreover, transfected cells of late passages did not reveal three-lineage differentiation potential (Suppl. Fig. S4a-c). Thus, the cells did not fulfil the minimal criteria for MSCs anymore^1^. Copy number variation (CNV) analysis confirmed that DNA of transfected cells (transfected^PL)^ at P4 with typical MSC morphology and P12 with epithelial-like morphology originated from the same donor, thereby ruling out the possibility that our results might be attributed to contamination with other cell lines (Fig. 3c). Notably, at late passage we observed a gain of large parts of the long arm of chromosome 1.

**Figure 3:**
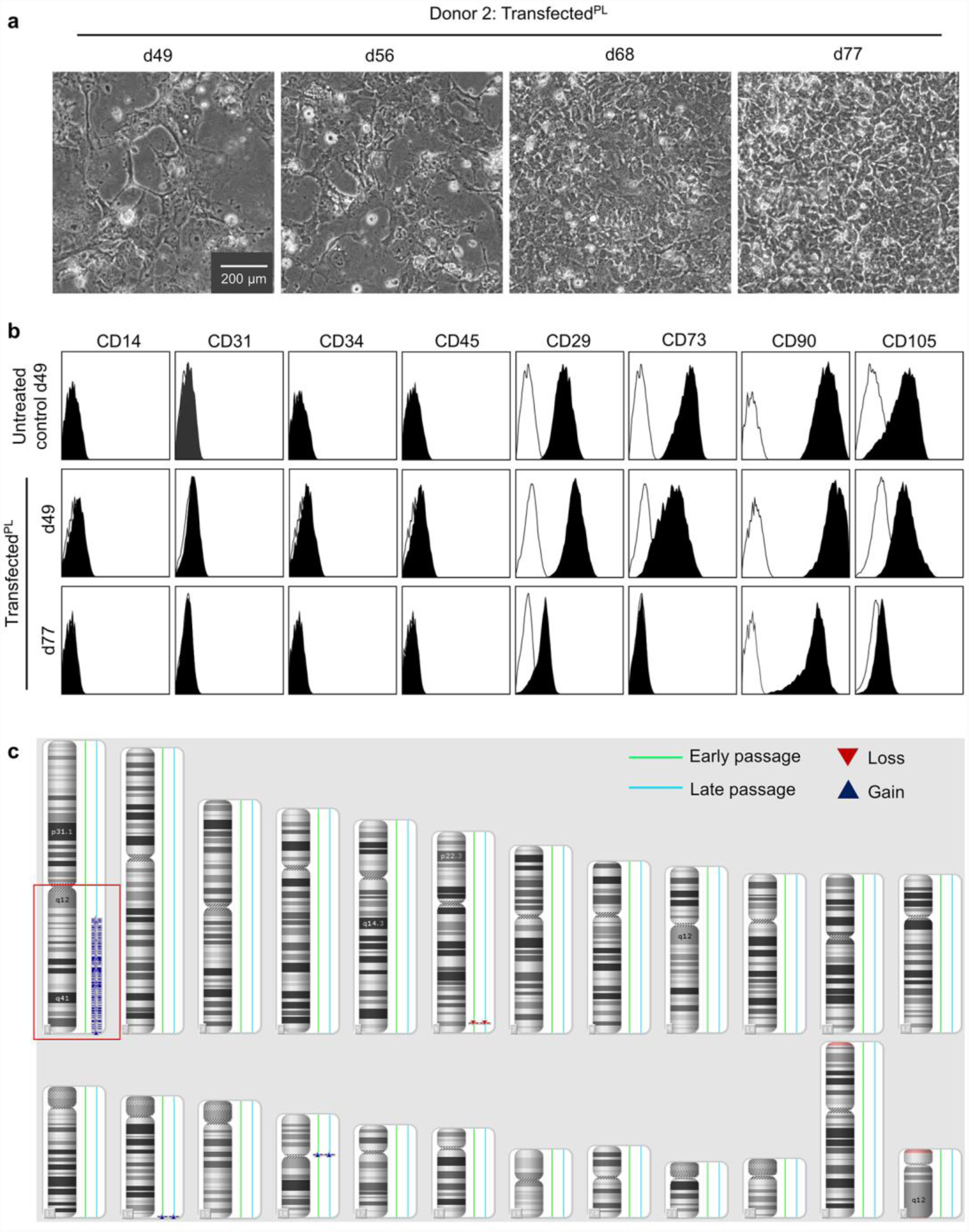
Transfection resulted in an aberrant subclone. (**a**) Phase contrast images of transfected cells of donor 2 (transfected^PL^) reveal pre-senescent MSC-like phenotype at d49 and then acquire an epithelial-like morphology. (**b**) Immunophenotypic analysis of untransfected (untreated control) and transfected cells (transfected^PL^) of donor 2 (10,000 events; log-scale). (**c**) Copy number variation (CNV) analysis with DNA of transfected cells (transfected^PL^) at passage 4 and passage 12 (only CNVs >200 kb with a mean marker distance of <5kb shown).

## Discussion

Partial reprogramming holds the perspective to counteract the aging process. However, our experiments did not provide any evidence that overexpression of pluripotency factors without reprogramming into iPSCs reverses senescence-associated modifications. This is in contrast to previous studies that described rejuvenating effects in mice with a doxycycline-inducible polycistronic cassette for OSKM expression^10,11^. Furthermore, Ocampo et al. used a human iPSC line with doxycycline-inducible OSKM factors to demonstrate that iPSC-derived fibroblasts have reduced γ-H2AX foci, a marker for DNA double-strand breaks, and restored H3K9 trimethylation levels upon short-term OSKM induction, while other markers for senescence were not addressed^10^. The discrepancy to our study might be attributed to the fact that we continuously passaged the cells every week, which reduces cell-cell interaction that is potentially crucial for early steps in reprogramming. In fact, we did not observe any iPSC colonies in the transfected^PL-i^ and the transfected^PL-i(MEF)-PL^ conditions, which usually give rise to iPSC colonies if passaging is avoided^19^. Another important difference is that the above-mentioned studies used a doxycycline-inducible expression system for OSKM factors that facilitates better titration and more homogeneous expression of pluripotency factors. Ocampo et al.^10^ used cyclic expression (2 days doxycycline, 5 days break, 35 cycles) and Guo et al.^11^ continuously induced OSKM expression for three weeks. We and others have previously demonstrated that the load of episomal plasmids rapidly declines within the first week and that they are no more detectable after several passages^12,19^. Yet, expression of pluripotency factors by episomal plasmids is notoriously heterogeneous and can hardly be controlled. Future experiments might therefore consider alternative strategies for overexpression of reprogramming factors, such as mRNA, retroviral, or lentiviral transfection^20^.

More importantly, we observed that transfection of one sample resulted in phenotypic and functional changes of MSCs with apparent immortalization. The simultaneous outgrowth of those aberrant cells in three culture conditions (PL, PL-I, and PL-i-PL) indicates that the genetic abnormality was acquired during transfection or early thereafter. After the initial 14 days of culture in hPL-medium the progeny of this cell was distributed to different culture conditions that were analysed in parallel. Spontaneous transformation of primary MSCs was previously attributed to contaminations with other cell lines^21^, while it seems to occur very rarely in primary MSCs under conventional culture conditions^22,23^. On the other hand, it has been demonstrated that premature termination of reprogramming after seven days by removal of doxycycline in the murine OSKM system resulted in tumour development resembling Wilms tumour^24^. Thus, the transformation event observed in our study might either be associated with epigenetic conversion due to partial reprogramming or with the genetic abnormalities that might be caused by the electroporation procedure. Either way, the results of our study exemplify that there is a risk with partial reprogramming that hinders using this approach for clinical therapy.

## Methods

### Cell Culture

Mesenchymal stromal cells were isolated from the femoral bone marrow of different donors after orthopaedic surgery as described before^25^. All samples were taken after informed and written consent and the study was approved by the ethics committee of RWTH Aachen University Medical School (permit number: EK300/13). Culture medium consisted of Dulbecco’s modified Eagle medium (DMEM, 1 gL^−1^ glucose; PAA, Pasching, Austria) supplemented with 1% penicillin/streptomycin (PAA), 1% L-glutamine (200 mM; Thermo Scientific, MA, USA); 10% pooled human platelet lysate that was generated as described before^26^, and 0.1% heparin (5000 lU mL^−1^, Ratiopharm, Ulm, Germany). Cells were cultured at 37 °C, in an atmosphere containing 5% CO_2_, and passaged by trypsinization every week with a reseeding density of 10,000 cells cm^−2^. Cells were counted using a Neubauer counting chamber and cumulative population doublings were calculated for every passage as described before^27^.

### Overexpression of pluripotency factors

MSCs were transfected with pluripotency factors using episomal plasmids as described before^19^. In brief, MSCs at passage three or six were trypsinized and 10^6^ cells were resuspended in 100 µL of R buffer (Life Technologies, Carlsbad, CA, USA). Subsequently, 1 mg/100 µL of each of the episomal plasmids (OCT3/4, siRNA for *p53*: pCXLE-hOCT3/4-shp53-F; SOX2, KLF4: pCXLE-hSK; L-MYC, LIN28: pCXLE-hUL; GFP: pCXLE-EGFP; and puromycin resistance [if indicated in the text]: pMSCV-puro; all Addgene, MA, USA) was added to the cell suspension. Transfection was carried out with the NEON transfection system according to the manufacturer’s instructions (Life Technologies; 1,700 V, 20 ms, 1 pulse). Subsequently, cells were seeded onto 6-well plates containing antibiotic-free hPL-medium in a density of 10,000 cells cm^−2^. The next day, medium was either exchanged for standard hPL-medium with or without puromycin dihydrochloride (Thermo Scientific) as indicated in the text. For reprogramming into iPSCs, the transfected MSCs were cultured on irradiated feeder layers of mouse embryonic fibroblasts (MEFs)^19^ in iPSC-medium consisting of Knockout DMEM with 20% Knockout serum replacement, 100 U/mL penicillin/streptomycin, 2 mM L-glutamine, 0.1 mM ß-mercaptoethanol (all Thermo Scientific) and 10 ng/mL basic fibroblast growth factor (bFGF; PeproTech, NJ, USA).

### Epigenetic signatures

Pluripotency was validated with the Epi-Pluri-Score based on DNA methylation at three CpGs^13^. In brief, genomic DNA was isolated using the NucleoSpin^®^ Tissue Kit (MACHEREY-NAGEL, Düren, Germany), bisulfite converted using the EZ DNA Methylation™ Kit (Zymo Research, Irvine, USA), and analysed by pyrosequencing on a PyroMark Q96 ID System (Qiagen, Hilden, Germany) at the three CpGs associated with the genes *POU5F1* (Oct4), *ANKRD46*, and *C14orf115*^13^. Alternatively, the state of cellular senescence was estimated with the Epigenetic-Senescence-Signature based on six senescence-associated CpGs related to the genes *GRM7*, *CASP14*, *CASR*, *SELP*, *PRAMEF2,* and *KRTAP13-3*, as described previously^17^. The DNA methylation values were then implemented into linear regression models to predict passage numbers^18^.

### Senescence-associated beta-galactosidase (SA-ß-gal) staining

This assay was performed using the Senescence Detection Kit ab65351 (Abcam, Cambridge, UK) according to the manufacturer’s instructions and analysed with an Evos FL Cell Imaging System (Thermo Scientific).

### Quantitative real-time reverse transcription PCR (RT-PCR)

Total RNA was isolated from cell pellets using the Nucleo Spin^®^ RNA Plus Kit (MACHEREY-NAGEL, Düren, Germany) and reverse transcribed into cDNA using the High-Capacity cDNA Reverse Transcription Kit (Applied Biosystems, CA, USA). Semi-quantitative RT-PCR was performed in duplicates on an Applied Biosystems StepOnePlus device using the Fast SYBR^®^ Green Master Mix (Applied Biosystems). Specific primers for human *CDKN2A* (*p16^Ink4a^)* were synthesized by Metabion International AG, Planegg, Germany (FW: 5’-CAACGCACCGAATAGTTACG-3’; RV: 5’-AGCACCACCAGCGTGTC-3’). *GAPDH* (FW: 5’- GAAGGTGAAGGTCGGAGTC-3’; RV: 5’-GAAGATGGTGATGGGATTTC-3’) was used as reference.

### Immunophenotypic analysis

Surface marker expression was analysed with a FACS Canto II (BD Biosciences, NJ, USA). The following antibodies were used for immunophenotypic analysis: CD14-allophycocyanin (APC; clone M5E2), CD29-phycoerythrin (PE; clone MAR4), CD31-PE (clone WM59), CD34-APC (clone 581), CD45-APC (clone HI30), CD73-PE (clone AD2), CD90-APC (clone 5E10; all from BD Biosciences) and CD105-fluorescein isothiocyanate (FITC; clone MEM-226; ImmunoTools, Friesoythe, Germany).

### *In vitro* differentiation of MSCs

Adipogenic, osteogenic, and chondrogenic differentiation of MSCs was induced as described before^28^. Briefly, cells were cultivated in the respective differentiation medium. After 21 days, fat droplet formation upon adipogenic differentiation was analysed by staining with BODIPY (4,4-difluoro-1,2,5,7,8-pentamethyl-4-bora-3a,4a-diaza-s-indacene; Invitrogen, CA, USA) and counter-staining with DAPI (4′,6-diamidin-2-phenylindol; Molecular Probes, CA, USA). Osteogenic differentiation was analysed by staining of alkaline phosphatase with NBT (nitro-blue tetrazolium chloride) and BCIP (5-bromo-4-chloro-30-indolyphosphate p-toluidine salt; Sigma Aldrich, MO, USA). Chondrogenic differentiation was assessed with Alcian Blue staining in combination with Periodic acid-Schiff (PAS).

### Copy number variation (CNV) analysis

Genomic DNA of transfected cells (transfected^PL^) at passage 4 and passage 12 was isolated as described above. For CNV comparison, the CytoScan^®^ HD Array (Affymetrix, CA, USA) was applied. Only CNVs >200 kb with a mean marker distance of <5kb were considered.

### Statistics

All experiments were performed with three independent biological replicas and results are presented as mean ± standard deviation (SD). Statistical significance was estimated by two-tailed paired Student’s t-test.

## Acknowledgements

We thank all the patients of this study for their collaborative participation. This work was supported by the Else Kröner-Fresenius-Stiftung (2014_A193), by the Deutsche Forschungsgemeinschaft (DFG; WA 1706/8-1 and WA1706/11-1), by the Interdisciplinary Centre for Clinical Research (IZKF; O3-3) within the faculty of Medicine at the RWTH Aachen University, and by the Flow Cytometry Facility, a core facility within IZKF Aachen.

## Author Contributions

C.G., R.G., and W.W. designed the study. C.G. and R.G. performed experiments, analysed and formatted the data. T.E. performed CNV analysis. C.G., R.G., and W.W. wrote the first draft of the manuscript and all authors read, edited, and approved the final manuscript.

## Additional Information

### Supplementary Information

Supplementary information is one PDF file with figures S1 to S4.

